# Towards a Brighter Constellation: Multi-Organ Neuroimaging of Neural and Vascular Dynamics in the Spinal Cord and Brain

**DOI:** 10.1101/2023.12.25.573323

**Authors:** Dmitrijs Celinskis, Christopher J. Black, Jeremy Murphy, Adriel Barrios-Anderson, Nina Friedman, Nathan C. Shaner, Carl Saab, Manuel Gomez-Ramirez, Diane Lipscombe, David A. Borton, Christopher I. Moore

## Abstract

**Significance:** Pain is comprised of a complex interaction between motor action and somatosensation that is dependent on dynamic interactions between the brain and spinal cord. This makes understanding pain particularly challenging as it involves rich interactions between many circuits (e.g., neural and vascular) and signaling cascades throughout the body. As such, experimentation on a single region may lead to an incomplete and potentially incorrect understanding of crucial underlying mechanisms.

**Aim:** Here, we aimed to develop and validate new tools to enable detailed and extended observation of neural and vascular activity in the brain and spinal cord. The first key set of innovations were targeted to developing novel imaging hardware that addresses the many challenges of multi-site imaging. The second key set of innovations were targeted to enabling bioluminescent imaging, as this approach can address limitations of fluorescent microscopy including photobleaching, phototoxicity and decreased resolution due to scattering of excitation signals.

**Approach:** We designed 3D-printed brain and spinal cord implants to enable effective surgical implantations and optical access with wearable miniscopes or an open window (e.g., for one-or two-photon microscopy or optogenetic stimulation). We also tested the viability for bioluminescent imaging, and developed a novel modified miniscope optimized for these signals (BLmini).

**Results:** Here, we describe novel ‘universal’ implants for acute and chronic simultaneous brain-spinal cord imaging and optical stimulation. We further describe successful imaging of bioluminescent signals in both foci, and a new miniscope, the ‘BLmini,’ which has reduced weight, cost and form-factor relative to standard wearable miniscopes.

**Conclusions:** The combination of 3D printed implants, advanced imaging tools, and bioluminescence imaging techniques offers a new coalition of methods for understanding spinal cord-brain interactions. This work has the potential for use in future research into neuropathic pain and other sensory disorders and motor behavior.

## 1 Introduction

Emergent biological phenomena, such as mammalian behavior, inherently depend on multiple computations conducted by distinct subsystems and the real-time interactions between them. While iterative study of subsystems in isolation can be highly beneficial, conjoint dynamics across systems, which could reflect distributed shared computations or essential interactive updating of distinct computations, are also essential to comprehending system function. As such, understanding these complex interdependencies also requires simultaneously recording biological activity across multiple organs.

Pain is a canonical example of a complex emergent phenomenon dependent on multiple subsystems. This clinically essential problem that is still insufficiently understood and addressed. Painful sensations evolved from nociceptive signaling of peripheral etiology, and implicate a wide array of chemical interactions and cell types (**Fig. 1** lists a subset of implicated signals). These signals navigate from skin to spinal cord, then ascend to the brain. While this classic ‘feedforward’ pathway description is intuitive, re-entrant feedback loops exist at all levels that impact pain sensation, including local reflex loops,^8,9^ descending projections^10^ and motor behavioral modification.^8,11,12^

**Fig. 1.**
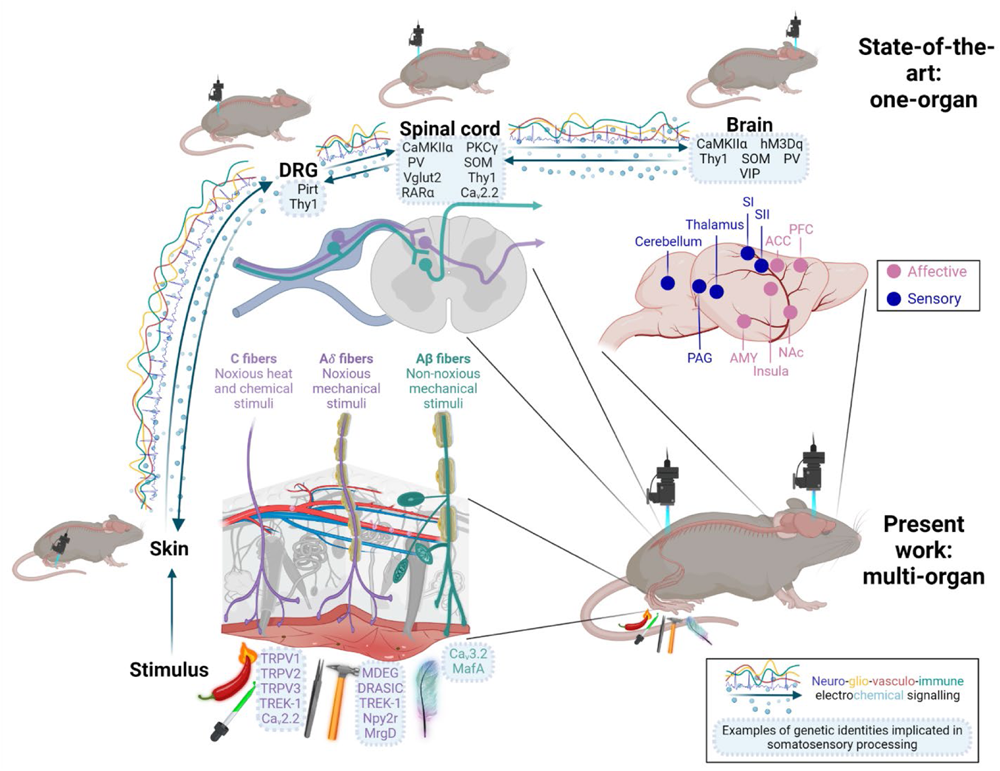
A Key Use Case for Multi-Organ Imaging: Studying Mechanisms of Nociception and Pain. Peripheral nociceptive input that drives pain engages a wide range of dynamic body systems. Integrated activity spans multiple organs, including skin, dorsal root ganglia (DRG), spinal cord and brain, and depends critically on a broad array of networks, including neural, vascular and immune signaling. State-of-the-art approaches for studying sensory information processing typically rely on in vivo imaging in one organ, and focuses on neurons. In the present work, we expand the in vivo methodological toolbox by developing procedures for multi-organ in vivo studies.

Direct imaging of activity and structure provides a powerful approach to capture the activity of specific cell types, and is typically essential to observe dynamics in the many non-neural systems that help create behavior and perceptual phenomena such as pain.^13^ Using genetically targeted fluorescent (FL) indicators, typical imaging studies target a single body area,^14–16^ including the brain,^14^ spinal cord,^15,17^ dorsal root ganglion,^18^ skin,^19,20^ heart,^21^ or intestinal tissue.^22^ Single area approaches necessitate using independent animals to study each site, and miss the potentially crucial interdependence between different body parts in generating phenomena. Further, this approach also leads to an increased number of animals needed for studies, leading to greater costs, unexplained experimental variability and higher animal numbers. Imaging methods such as functional magnetic resonance imaging (fMRI) that provide whole-brain and spinal cord access with the ability to resolve and stimulate single brain regions^23,24^ and brain-spinal cord at the same time^25^ have been transformative to our understanding of multi-subsystem interdependencies in behavior. However, fMRI lacks the single-neuron resolution, temporal precision, and genetic specificity possible with optical approaches.

While all these factors suggest the need for multi-organ optical access with single-cell precision, there are several challenges to achieving this goal in mice. A significant hurdle to making multi-organ microscopy standard in live animals is the increased surgical complexity. Implantation of cranial and vertebral implants is typically done under anesthesia, and combining both procedures in a single animal can require longer loss of consciousness and increased risk of anesthetic exposure. As one example, isoflurane, in standard use for such implantations, shows cumulative effects of exposure^26^ and can impact microstructural organization^27^ and protein phosphorylation in the brain^28^, cause postoperative cognitive dysfunction,^29^ increase susceptibility to anxiety-like behaviors,^30^ and modulate cardiovascular function.^31^ Importantly, exposure to isoflurane has also been reported to affect calcium dynamics in the spinal cord,^15^ stimulate peripheral sensory nerves by sensitizing transient receptor potential V1 (Trpv1) receptors,^32^ inhibit neuronal CaV3.2 calcium channels,^33^ and increase the susceptibility to chronic pain by causing alterations in dorsal spinal cord and dorsal root ganglion.^34^ Given these basic challenges, a key requirement for multi-site imaging is the design of surgical procedures that minimize duration, and that allow awake imaging if possible.

Further, the ability of animals to tolerate multiple window implants while maintaining health is also an open question. In addition to the potential immunological load created by multiple implants, they also create unique mechanical challenges, including steric hindrance between sites, and up to a doubling in the weight the animal must tolerate. In a small creature such as a mouse, miniaturization of head implants, for example, needs to fall under ∼2g to avoid impeding behavior and normal motor function.^35^ Also, brain and spinal cord surgeries in mice require fine motor dexterity and nuanced microsurgical skills. Hence, availability of the neurosurgery personnel, ability to invest time into learning new surgical procedures and optimizing surgical training protocols to reduce training animal use are additional important considerations when implementing a novel surgical procedure as we present in this report.

FL imaging has been a remarkable, high-signal approach for tracking neural activity during behavior.^36–38^ However, FL imaging is limited by its relatively high noise, created by factors such as tissue autofluorescence.^39,40^ Dependence on excitation light also adds noise to small-target imaging in turbid media, as photons sent into tissue to elicit a FL scatter on the inbound path, creating image blurring and constraining imaging depth (**Fig. 2**). Further, continuous application of external light, as typically employed with miniature microscopes (‘miniscope’) imaging, causes photobleaching,^41^ phototoxicity,^42^ and can adversely affect cellular capacitance^42,43^ and mitochondrial function.^44^ Additionally, FL miniscopes have been a great addition to the armamentarium of new tools for studying brain activity,^45–49^ benefitting from low weight (2 grams)^50^ and effective optics. However, using this new gold standard for multi-site imaging is a challenge, given the cumulative weight of multiple scopes, and this size issue grows with the need to increase simultaneously imaged colors or transition to wire-free scopes.

**Fig. 2.**
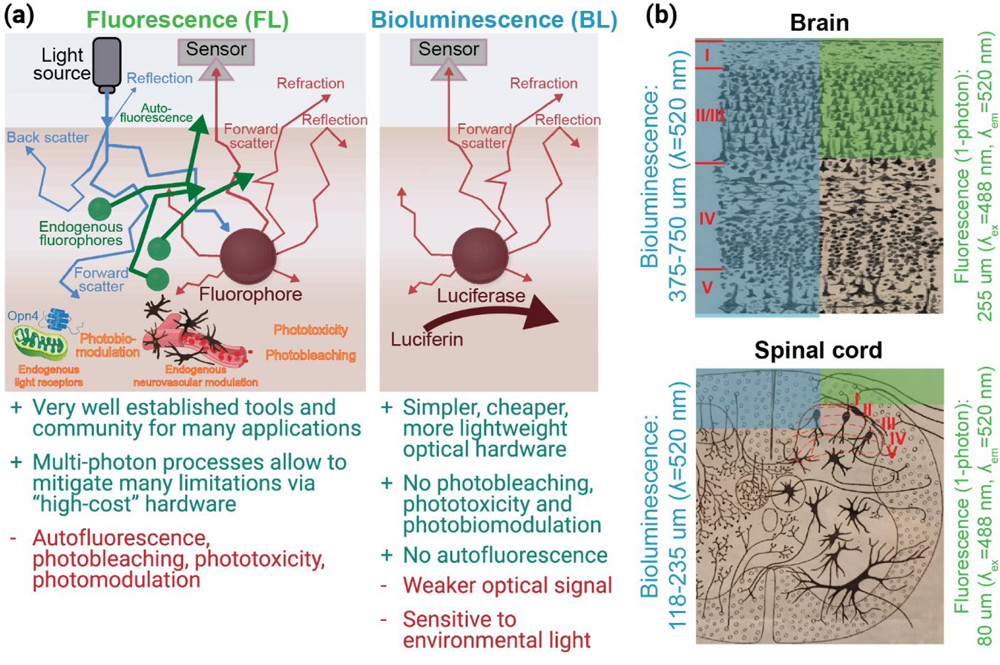
Imaging Bioluminescence (BL) Offers Advantages Over Fluorescence (FL), Including Decreased Scattering Artifacts and Simplified Optics. **(a)** Comparative advantages of BL and FL imaging. **(b)** BL offers up to 3x potential increase in imaging depth of discrete emitters (e.g., cells) compared to FL. Mouse somatosensory cortical layers and spinal dorsal horn laminae marked in red indicate the approximate deepest cortical layers^1–3^ and laminae^2,4^ that can be imaged in an adult mouse via 1-photon FL and BL approaches. Imaging depths were determined from.^5^ The estimated increase in imaging depth is especially beneficial for spinal cord imaging because it is significantly constrained by photon scattering properties of myelin. The brain background image in (b) is from^6^ and is in the public domain, the spinal cord image in (b) is from,^7^ licensed under Public Domain Mark 1.0.

These issues with FL imaging may be addressed with bioluminescent (BL) indicators. BL indicators can provide robust calcium signals with relatively high spatio-temporal resolution.^51–55^ Because BL signal production depends on a chemical interaction at the site of the indicator and not incoming light, it removes the need for excitation light. The noise due to scattering is essentially reduced by 50% (**Fig. 2**), photodamage is removed almost entirely as a concern, and the removal of the excitation light further reduces the overall device weight. Further, for studies of multi-site interactions during behavior, distinct BL indicators (e.g., emitting different colored signals) can be targeted to different subsystems throughout the body, and monitored without the need for implants.

Here, we took several steps towards a complete system for imaging neural and vascular dynamics in the brain and spinal cord with cellular resolution within the same mouse. We have developed custom implants that simplify and make brain-spinal cord procedures more accessible by reducing implant cost and fabrication complexity, and by streamlining customization through 3D-printing. Our custom implant offers more versatility compared to prior implants, adapting to a wider range of use cases: brain or spinal cord, restrained or free behavior, wearable 1-photon or benchtop 1-and 2-photon imaging, and acute or chronic imaging for up to 230 days post-surgery. We show how to (i) fabricate versatile brain-spinal cord window implants, (ii) perform surgeries for brain and spinal cord within the same animal, (iii) habituate multi-implant animals for behavior, (iv) image anesthetized and awake animals simultaneously or sequentially across the brain-spinal cord, and (v) optically and mechanically modulate activity across these areas.

We take several steps to enabling BL imaging that can overcome key constraints. We demonstrate that robust BL signals can be imaged in the brain. Further, we refined the design of UCLA miniscopes to improve signal sensitivity, reduce miniscope’s weight and simplify the assembly of hardware.

## 2 Materials and Methods

### 2.1 Animal Subjects

All procedures were approved by the Institutional Animal Care and Use Committee (IACUC) at Brown University. Mouse strains used in this study were purchased from the Jackson Laboratory and bred in-house: C57BL/6 wild-type mice, SOMcre x tdTomato x Ai148 (TIGRE-Ins-TRE2-LSL-GCaMP6f; JAX# 030328), Trpv1^Chr2EYFP+/−^ (Trpv1^Cre+/+^ is JAX#: 017769 and lox-STOP-lox-ChR2-EYFP is JAX#: 012569), Cacna1h^Chr2EYFP+/−^ (Cacna1h^Chr2EYFP+/−^ line were developed in the Lipscombe laboratory by mating Cacna1h^Cre+/+^ with ChR2-EYFP^+/+;^ Cacna1h^Cre+/+^ is also known as CaV3.2^Cre+/+^; the Cacna1h^Cre+/+^ mice used for this project were inbred 13 times). Mice were group-housed until the implant surgery and individually housed after the surgery. Mice were provided with bedding and nesting materials, and maintained on a 12 hour light-dark cycle in a temperature and humidity controlled environment.

### 2.2 Implant Fabrication

For ease of reference, we refer to our **U**niversal format 3D-**Pr**inted **Im**plant as the UPRIM. All the implants tested for this work were 3D-printed via stereolithography (SLA). The 3D-printing photopolymer resins tested in this work included: (i) Accura Xtreme White 200, (ii) Accura SL 7820 in black color, (iii) Accura Xtreme in gray color, (iv) Accura 60, (v) Accura ClearVue, (vi) Dental SG (Formlabs, FLDGOR01), (vii) Tough 1500_v1 (Formlabs, FLTO1501). We found both Dental SG and Tough 1500_v1 to be well suited for fabricating the spinal bars and implant chambers (**Fig. 3**). Dental SG offers better biocompatibility than Tough 1500_v1. However, based on the qualitative observations, we did not observe any differences between two the implants from these resins in terms of post-surgical animal behavior and tissue quality around the implant. The important considerations to keep in mind is the relatively high translucency and brittleness of the implants made from Dental SG resin. Transparency can be important in the optogenetic studies where it might be undesirable to confound the experimental results because of the endogenous effects resulting from the light exposure outside of the implanted window. Brittleness in our hands was a concern only when trying to thread the implant chamber directly. In contrast with the Tough 1500_v1, direct threading of Dental SG implants did not allow >1 screw-unscrew operation cycle. However, the issue of translucency can be addressed by painting the implant and the issue of brittleness can be resolved by gluing the commercially available nuts into the implant. All the results presented in this report were collected using UPRIMs fabricated from Tough 1500_v1 resin (**Fig. 3**).

**Fig. 3.**
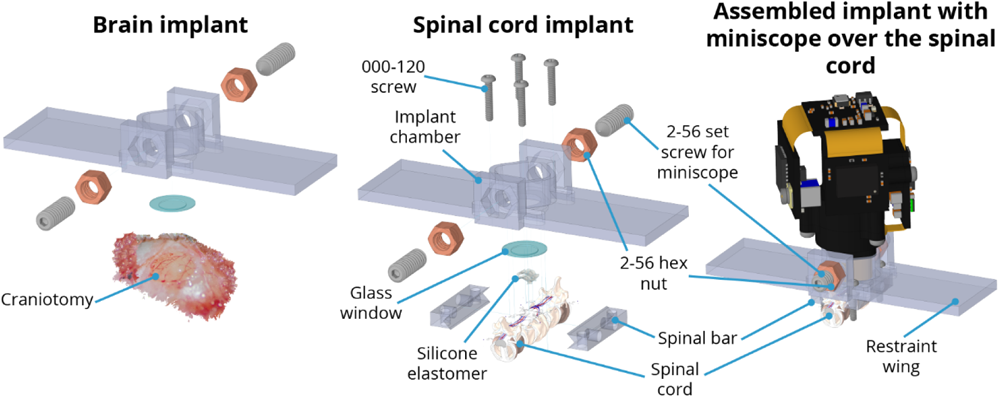
UPRIM Brain and Spinal Window Implants Consist of Off-the-Shelf Components and a 3D-Printed Implant Chamber. Identical UPRIM implant chambers can be used for brain and spinal imaging. The only material difference is the need for additional parts to attach 3D-printed chamber to the spinal cord. The specific design illustrated here is equipped with a mount for UCLA miniscope v4, but can also be readily adapted for any wearable microscope and is compatible with the benchtop microscopes with ≥ 5 mm objective working distance.

**Fig. 4.**
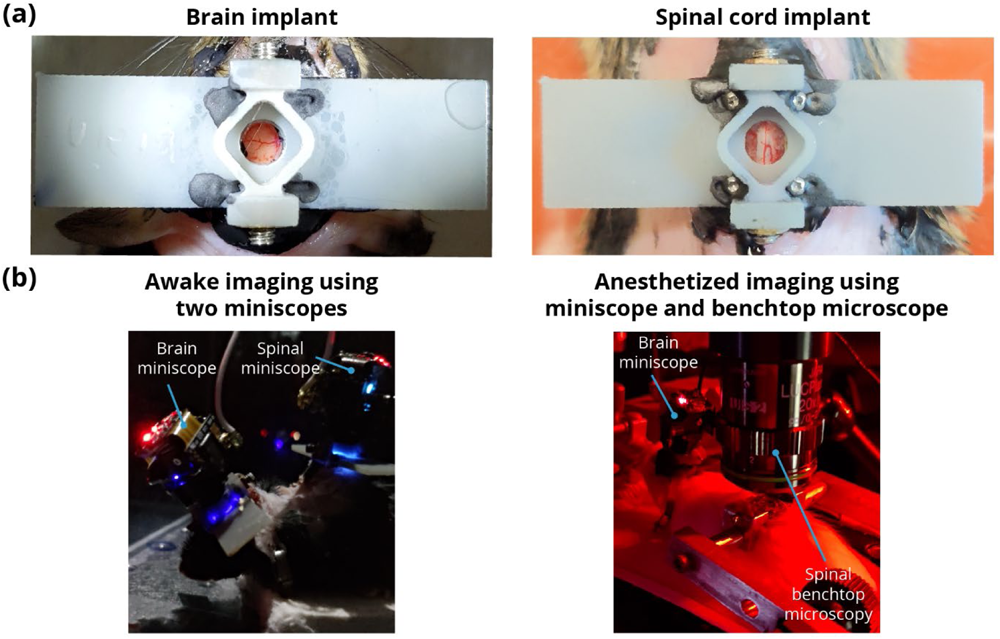
UPRIM Implants Can Be Used for Brain and Spinal Cord Imaging in Awake and Anesthetized Animals (a) Images of the UPRIM 3D-printed implant attached to the brain and the spinal cord. **(b)** Images of use of the implants for imaging through multiple miniscopes (UCLA V4) and a benchtop microscope.

The spinal cord bars and spinal cord chamber were fabricated from Tough 1500_v1 resin using an in-house 3D-printer (Form 3, early 2019 edition, FormLabs Inc.). The fabrication steps were as follows:

1. 3D-print the parts,
2. Wash parts in isopropyl alcohol (by hand or using Form Wash washer from FormLabs Inc.),
3. Perform only for the implant chamber: apply the UV-curing glue (electronic adhesive 123SBLUE, Norland Products Inc.) and insert the nuts (92736A101, McMaster-CARR Inc.),
4. Cure (using Form Cure, FormLabs Inc.),
5. Perform only for the implant chamber: after curing we usually apply additional small droplets of 5-minute epoxy (QUIK-CURE, bSi Inc.) at the interface between the nuts and the implant,
6. Remove the supports,
7. Sand off any residual marks left while removing the supports,
8. Thread and test the fitting of all the cylindrical holes:

a. spinal cord bars:

i. hand-drill the cavities for the bar holders and make sure they fit well on the surgical bar holders;
ii. manually tap the vertical holes and test the fitting of the 000-120 screws (90910A603, McMaster-CARR Inc.);
iii. implant chamber:
iv. test the fitting of the 2-56 ¼” set screws (90778A022, McMaster-CARR Inc.) into the glued nuts;
v. test the smoothness of the passage of 000-120 screws through the lateral vertical cavities used for threading the screws through the chamber into the spinal cord bars;
vi. test the fit of the miniscope into the implant chamber;
b. Autoclave before the surgery. At time of writing, outsourced high resolution stereolithographic 3D-printing from Accura 60 resin with quick clean finish is ∼$76/implant (e.g., at www.3dsystems.com), and 3D-printing cost becomes negligible if using in-house 3D-printing. In contrast, machined spinal cord implants cost on the order of $349/ implant for the individual implants or $171/implant when ordering in bulk (Neurotar Inc.). Further, our implant chambers are the first that allow imaging both in freely behaving animals with miniscopes and under restraint. Commonly available miniscope baseplates cost ≥$32/implant (miniscopeparts.com), but do not allow animal restraint and cannot be used for chronic spinal cord imaging.

### 2.3 Surgical Procedures

#### 2.3.1 Surgeries for brain AAV injections and window headpost implantation

We performed brain injections of adeno associated virus (AAV) and window headpost implantations as parts of a single procedure. Depending on the specifics of AAV injections, this surgical procedure takes on average 3-4 hours plus 1-2 hours of preparation time. All surgeries followed the aseptic techniques. We anesthetize mice using 1.5-2% isoflurane delivered in 100% oxygen gas and eliminated using vacuum suction. We protect the animal’s eyes using sterile ocular ointment and the animal’s body temperature is maintained stable using a heating blanket. We prepare the surgical site on the skin in a dedicated area away from the surgical sterile field. We inject the animal with intraperitoneal (IP) dexamethasone (0.2 ug/g), subcutaneous (SQ) meloxicam SR (4 ug/g) and local SQ lidocaine (3 ug/g). We then cut off the longer hair from the top of the animal’s head and apply a depilation cream (Nair hair-remover lotion, Church & Dwight Inc.) around the planned incision site. We massage the depilator cream into the hair and wash it off together with the detached hair using alcohol after 30 seconds. We then clean and sterilize the skin with a two-stage scrub of betadine and 70% ethanol, repeated three times. After cleaning and sanitizing the skin, we move the animal to the nearby surgical sterile site without interruption to the animal’s plane of anesthesia. We then stabilize the animal’s head using earbars. We use a 10 mm ring to pressure-mark the edges of the intended cut in the skin. We cut a circular piece of skin using scissors by following the previously marked trace. We then remove all the connective tissue remaining on the skull and cut the muscle pockets on both lateral sides of the skull. This step is most prone to result in multiple muscle bleeders, especially during the early stages of learning this procedure. During this step and throughout the rest of the surgery we use surgical absorbent gelfoam (Surgifoam 1972, Ferrosan Medical Devices) to control and stop any bleeders. We prepare gelfoam at the beginning of the surgery by applying saline to it to make the gelfoam soft while not free-floating in excessive amount of saline. Throughout the entire surgery we use only the gelfoam softened by saline. Next, using the surgical blade, we make densely distributed surface scrapes in the skull at the top and on the sides of the muscle pockets, and then etch the same skull surfaces using the dentin activator (Universal Dentin Activator Liquid S393, C&B-Metabond®, Parkell) for 10 seconds followed by a wash off with saline. Scraping and etching are important steps to ensure reliable long-term bonding of the dental cement to the skull and attachment of the implant. We then mark bregma and mark the location of somatosensory cortex hind-limb area (S1HL, AP = 0.5 mm and ML = 1.5 mm) relative to bregma. We use the ear punch (3mm disposable biopsy punch, 19D23, INTEGRA Inc.) by gently scraping the surface of the skull to mark out a 3 mm ring for drilling a craniotomy centered on S1HL. We use a Neoburr drill bit (FG ¼, Microcopy) for the initial drilling of the skull through 50-75% of the skull’s thickness. We drill through the rest of the skull in gradual pecking motion using a beveled cylinder drill bit (1812.8F, Microcopy). After completing every full drilling circle we apply saline to prevent overheating and flush away the bone chips. Once the craniotomy piece of the skull starts moving with brain pulsations, we remove this bone piece using forceps. We then clean the exposed brain from any blood, CSF and bone chips using a piece of gelfoam. From this point in the surgery, we always make sure the brain never dries out and is moist with saline. We maintain a good level of moisture both by regularly pouring saline over the craniotomy and keeping the craniotomy covered with gelfoam for longer periods of time. We then take a sterile insulin syringe (1144303, 0.5mL 28G, BD Insulin Syringe, Belton, Dickinson and Co.) and bend the tip of 31G needle by gently tapping the needle against a flat hard surface. This serves as a hook tool for removing the dura. We use such a syringe with the hook to make 2-3 cuts in dura for the upcoming viral injections. We let the brain stabilize after the durotomies while preparing for the viral injections.

Glass pipettes (5-000-1010, DRUMMOND WIRETROL, Drummond Scientific Co.) were pulled on a Model P-97 (Sutter Instrument Co.). We cut the tips using Vannas scissors under the microscope to the tip diameter of 25-50 um. We back fill the pipette with the mineral oil (O121-1, 148223, Fisher Chemical Inc.) and front fill the pipette with the virus using the injector (53311, Quintessential Stereotaxic Injector, Stoelting Co.). We injected Cacna1h^Chr2EYFP+/−^ animals with 500-750 nL of AAV2/9-CaMKIIa-GCaMP6f-P2A-nls-dTomato (Canadian Neurophotonics Platform Viral Vector Core Facility, RRID:SCR_016477) to the depths of 150-250 um. We injected Trpv1^Chr2EYFP+/−^ animals to the depths of 150-250 um with 500-1,500 nL of cocktail mixtures of AAV2/9-CaMKIIa-GCaMP6f-P2A-nls-dTomato and SYN-jRGeCO with GCaMP:jRGeCO proportion of 1:1. We mixed two AAVs and the green food dye (Fast Green FCF Solution, 26053-02, Electronic Microscopy Sciences Inc.) by releasing and withdrawing the mixture 2-3 times with a micropipette. We add green food dye to aid visualization of the flow of AAV during the injection. We insert the pipette into the brain using a stereotactic manual micromanipulator. We wait for 3 min after inserting the pipette to the targeted depth to allow tissue to stabilize and we inject the AAV at the rate of 60 nL/min. Following the injection, we wait for 5 minutes to allow the tissue to stabilize before withdrawing the pipette.

Once all 2-3 injections are complete, we cover the brain with the glass window. We make our glass windows by gluing together 3 mm (CS-3R, 64-0720, Warner Instruments Inc.) and 5 mm (CS-5R, 64-0700, Warner Instruments Inc.) circular coverslips using the optical adhesive (Optical adhesive 71, Norland Products Inc.). The 3 mm coverslip fits relatively snugly into the craniotomy, forming a tight seal against the drilled skull. This is important to minimize any risk of bone bleeders during the healing and to delay the overgrowth of craniotomy with dura and bone. We press the glass window into the brain using a basic custom pressure applicator with a diameter of 2 mm mounted on a stereotactic manipulator. Once the glass window is tightly pressed into craniotomy, we secure the window in place by applying a 3-component dental cement (C&B-Metabond®, Parkell) around the circumference of the window. Before applying the dental cement to the skull, we add a black paint to the 3-component dental cement mixture to reduce autofluorescence of the dental cement. Once we apply the dental cement around the circumference and the dental cement begins to harden, we withdraw the pressure applicator and start applying the dental cement throughout the rest of the skull. We also fill up with dental cement the muscle pockets that we make at the beginning of the surgery. Next, we cover the bottom of the implant with uncured dental cement mixture and place the implant on the skull centered over the cranial window. We then apply the dental cement mixture from all 4 sides of the implant to increase the security of its attachment to the skull. We also pour some dental cement mixture through the 4 holes atop of the implant that are used for attaching the screws during the spinal cord surgery below. Finally, we apply tissue adhesive (Gluture, Zoetis Inc.) throughout the perimeter of the dental cement to double secure the attachment of the dental cement mixture to the skull and the skin.

#### 2.3.2 Surgeries for Spinal Cord AAV Injections and Window Implantation

We perform spinal cord AAV injections and window implantations as part of a single surgical procedure. In our hands, the full procedure on average required 3-5 hours plus 1-2 hours of preparation time depending on the specific parameters of the AAV injections and specifics of any given animal. This procedure was developed by the right-handed surgeon and might require small modifications to achieve the optimal timing for the left-handed surgeons.

All surgeries follow the aseptic techniques. We anesthetize the animal using 1.5-2% isoflurane delivered in 100% USP-grade oxygen and exhausts collected using vacuum. We maintain the animal’s body temperature stable throughout the surgery using a heating blanket and protect the animal’s eyes using sterile ocular ointment. We prepare the animal in a non-sterile area before moving it to the sterile stereotactic frame. For the chronic spinal cord surgeries, we use the same stereotactic setup and injection tools as for the brain surgery, except we add a stack of breadboards (Thorlabs Inc.) with XYZ manipulators (60 mm x 60 mm footprint available from Amazon) needed for spinal cord manipulation. The spinal cord clamp holder posts used for all our surgeries were 220 mm in length. The original design by Farrar et al. 2012 included the spinal cord clamp holder posts 50 mm in length. We significantly increased the length of the implant holder posts to accommodate the dimensions of our stereotactic frame used for brain surgeries (Kopf Inc.). There were no particular benefits for using longer clamp holder posts. However, in our hands the use of XYZ manipulators rather than hand-manipulation like in Ref. ^17^ yielded finer control over the implant position and simplified the transfer of knowledge between the students by simplifying this complex procedure.

After moving the animal from the induction chamber to the prepping station, we inject the animal with IP dexamethasone (0.2 ug/g), SQ buprenorphine SR (0.75 ug/g), SQ glycopyrrolate (0.5 ug/g) and local SQ lidocaine (3 ug/g). We then generously shave the animal’s back using an electrical buzzer and apply a depilation cream (Nair hair-remover lotion, Church & Dwight Inc.) around the intended incision site. The area for depilation can be determined by finding the apex of the vertebral column’s curvature and the location of the last rib. The apex of the vertebral column curvature corresponds to the Th13 vertebrum.^56^ We massage the depilator cream thoroughly into the animal’s back using a cotton swab to avoid any unexposed hair and we remove the detached hair using a razor blade after 30-60 seconds. We thoroughly wash off any remaining depilator cream and detached hair using alcohol pads. We then clean and sterilize the exposed skin with a two-stage scrub of betadine and 70% ethanol, repeated three times. After this, we move the animal to the surgical stereotax for the sterile parts of the procedure.

We begin by aligning the animal’s back with the XYZ manipulators to make sure the spinal cord clamps mounted on the custom clamp holder posts can reach the XYZ position of the spinal cord. We then prepare the surgical gelfoam by soaking it in sterile saline. We proceed by identifying the location for the skin incision by palpating the ribs around the apex of the vertebral curvature. Typically palpation involves active sweeping of the back with the fingers, as well as keeping the fingers in place around the suspected last rib to feel the motion of the last rib while breathing. The last rib emanates from the rostral end of the Th13 vertebrum.^56^ For the imaging of the spinal cord at the vertebral level L1, we usually aim to have the skin incision that starts about 1-3 mm rostrally relative to the entry point of the last rib, covers the width of the implant (10 mm) and extends 1-3 mm caudally (i.e. 12-16 mm total). As was described in Ref. ^17^, it is possible to obtain a good quality spinal cord window even with a significantly larger skin incision. However, it is important to minimize any tissue damage and trying to tailor the incision size towards the implant’s size does not have a significant impact on the duration or complexity of the surgery. For the imaging of the spinal cord segments under the L2 vertebrum, the skin incision should be moved caudally by 2.5-3.0 mm. Following the skin incision, we repeat the palpation of the last rib this time using the forceps (Fine Instruments Inc.) to locate the rostral end of Th13 more precisely. Once we locate the root of the last rib, we incise the soft tissue overlying the vertebra. The length of the cut is equivalent to the width of the implant. We begin by making a straight cut on the right side as close to the midline as possible while not cutting through the spinous processes. We then turn the surgical blade by more than 70 deg to orient it close to the horizontal plane and cut laterally towards the right to dissect away the overlying soft tissue. After reaching the transverse processes, we reorient the blade vertically and dissect away the tissue off the vertebral bodies not deeper than the thickness of the spinal cord bars (1.5 mm) to offer enough space for fitting the spinal cord bars under the spinal cord processes flush against the vertebral body. Similar to Farrar et al. 2012, the implant bars merge together 3 vertebrae. Hence, we clean up the lateral surfaces of 3 vertebral bodies, as well as cut off the tendons attached to all 3 vertebral bodies to minimize the motion during post-surgical imaging. After cleaning the lateral sides of vertebral bodies, we clamp the vertebral column from both sides with the spinal cord bars. We align the center of the bars with the approximate center of planned laminectomy over the middle vertebrum. We apply a generous amount of pressure to keep the spinal cord in place during the surgery and to promote strong chronic attachment via pressure after the surgery. Next we partially trim away the spinous and transverse processes to maximize the flatness of the implant chamber against the vertebral bodies. The processes are not removed entirely, rather they are flattened by cutting away the superficial parts of the processes while keeping as big parts of the processes as possible to minimize tissue damage and promote strong attachment of the spinal cord bars. In our practice, it is best to cut the processes as early as possible because this step is prone to bone bleeding if done improperly and by cutting the processes early, the tissue has a longer time for coagulation before further cuts must be made. Next, we clean up the intervertebral spaces from all the connective tissue to have a clear tactile and visual feedback on the extents of the middle vertebrum. Once we know the extents of the vertebrum, we perform a laminectomy using a drill actuated via compressed nitrogen (both diamond burr and ball drill work well for this step). We sequentially drill longitudinally as far laterally as possible first on one side, then on the other and repeat switching sides after every pass, removing thin layers of bone on every pass until the bone becomes thin enough to be pulled up without having to apply a significant amount of force. Between every pass of the drill we apply the sterile saline to protect the spinal cord from drying.

After the laminectomy, we perform the viral injections following the same steps as for the brain injections. The main difference between the brain and spinal cord viral injections in our experience is that the spinal cord meninges are much harder to pierce through compared to the brain meninges. Hence, we performed durotomies for every spinal cord injection. The examples of pinching responses in Fig. 16 involved injections of an AAV2/9-CaMKIIa-GCaMP6f-P2A-nls-dTomato (Canadian Neurophotonics Platform Viral Vector Core Facility, RRID:SCR_016477) virus at 60 nL/min. We administered the AAV via two injections on the right side of the spinal cord at 150 um and 250 um depths and one injection on the left side at 200 um.

#### 2.3.3 Multi-Organ Surgeries as a Single or Two Separate Procedures

When performing 2 independent surgeries within the same animal, each procedure is performed exactly as outlined above. The main important factor to consider is that the animal requires at least 3 days to recover after a brain or spinal cord surgery. We performed the second surgery 7 days after the first surgery out of an abundance of caution. Further, we typically performed brain surgery first followed by the spinal cord surgery. This choice was based on our experience with maintaining the good quality and responsiveness of surgerized neocortex across months without significant challenges with good consistency between the animals. Hence, the risk of the cranial window to decrease in quality while waiting for the spinal cord window (e.g. waiting for healing or viral expression) is lower than the risk or the amount of unavoidable quality change for the spinal cord window.

In the case of both brain and spinal cord surgeries performed as part of a single procedure, we follow the same surgical steps as outlined above except in all surgical cases the animals did not receive any viral injections in either of the sites. This was due to the significant surgical duration and the impact on the animal’s health when trying to incorporate AAV injections into the surgery. The duration of a combined surgery in our implementation is at least 6 hours. We typically would start with a spinal cord implant first and then would proceed with the brain implant. This choice is based on the fact that the spinal cord surgery is more prone to failure and if the spinal cord surgery is done following the brain surgery, the resources invested into brain surgery would be wasted.

As part of the development of surgical methodology it was important to ensure that the surgical developments presented in this report are transferable to other scientists. The transferability of principles described here was validated by training a medical student on brain-spine surgeries. The transfer of knowledge could be efficiently performed following the knowledge and materials developed throughout this work and trained student was able to successfully surgerize the animal that was used to generate the data presented in **Fig. 10a**.

### 2.4 Fluorescence Imaging

#### 2.4.1 One-photon Imaging Using Electron Multiplying Gain CCD (emCCD) Camera

The results shown in **Fig. 10b** were imaged using an emCCD camera. For imaging calcium using a green FL protein based genetically encoded calcium indicator (GCaMP) and the blood vessels labeled with the fluorescein isothiocyanate (FITC) dextran (**Fig. 11 and 13c**) the emCCD camera was equipped with GFP filter cube (DFM1 with GFP excitation and emission filters and dichroic, ThorLabs) and an excitation LED (M470L3, ThorLabs Inc.).

#### 2.4.2 Two-photon Imaging

All the 2-photon imaging was done using a custom microscope assembled by Bruker Inc. Vascular imaging data shown in **Fig. 11** were collected using a 960 nm excitation laser wavelength with galvo scanning and 16x water immersion objective (0.80W, DIC N2, Nikon LWD). Brain calcium imaging data shown in Fig. 18 was collected using a 980 nm excitation laser wavelength with galvo scanning and 10x objective (HC PL APO 0.40NA, 506511, Leica Inc.).

#### 2.4.3 Imaging Using Miniscopes

All the miniscope imaging was performed using UCLA miniscopes v4.0, v4.40 and v4.41. Miniscope v4.0 was generously provided by UCLA, v4.41 was purchased from OpenEphys and v4.40 was purchased from OpenEphys. v4.40 miniscope from LabMaker offered the smallest amount of bleed-through excitation light noise and best imaging quality. Miniscopes v4.40 and v4.41 had the lens configuration #1 (combination of lenses #45-089, #45-089 and #45-691 from Edmund Optics with working distance 675 um +/-250 um and FOV of 1 mm x 1 mm) [250] and was used for brain and skin imaging. Miniscope v4.0 had the lens configuration #3 (combination of lenses #45-090, #45-090 and #45-691 from Edmund Optics with working distance 2010 um +/-400 um and FOV of 1.3 mm x 1.4 mm) and was used for spinal cord imaging. We tested the lens configurations #1 and #2 (working distance for configuration #2 is 975 um +/- 340 um) for spinal cord imaging in multiple animals, but the working distance was too short for imaging using the window design and viral injection depth combinations used in this work. In 90% of the cases all lens configurations allowed us to at least reliably image the cortical blood vessels.

For the imaging under anesthesia, the miniscope was attached to the implant chamber using 1 implant screw and for the awake imaging using 2 screws. We also imaged anesthetized animals implanted with the stainless-steel headposts and cranial windows not designed for the attachment of the miniscope. During these imaging sessions the miniscope was mounted in a 3D-printed holder and positioned over the cortex using micromanipulators.

#### 2.4.4 Vascular Imaging Using Dextrans

Blood vessel imaging reported in **Fig. 11** relied on labeling the blood vessels using injectable FL dextran dyes. Dextran was delivered into the animal’s body either via tail vein injection. Dextran was injected by hand using a syringe. The dextran used in this study were as follows: FITC 10 kDa (D1821, ThermoFisher Scientific Inc.) delivered as a 30 uL of dextran solution mixed with 70 uL of sterile saline.

#### 2.4.5 Behavior for Fluorescence Imaging Experiments

The awake imaging results reported in **Fig. 10c** required the animal to follow the 7-step behavior procedure summarized in **Fig. 5**. The steps were as follows:

1. Handling

a. Handling helps to improve an animal’s collaborativeness throughout the experiments. Given how sensitive the spinal cord preparations can be depending on the study and surgeon’s technique, as well as the complexity of multisite surgical procedures, it can be beneficial to assess an animal’s collaborativeness ahead of the first surgery. Animals that within 3-5 days do not become significantly better at collaborating with the researcher and do not increase in comfort of being handled, it can be beneficial for animal’s health and the experiment to assign a given animal to the acute track or invest additional time to increase the chances for a given animal’s successful participation.
b. Handling helps to maximize the animal’s comfort with the experiment, minimize the risk of tissue damage, increase the experimental duration and overall, improve the quality of data.
c. Increasing the animal’s comfort with being handled by the researcher is especially critical for maintaining the longevity of the spinal cord implants. Even miniscope imaging requires restraining or anesthetizing the animal for 2-5 minutes to attach the miniscope. Hence, by the first time of needing to attach a miniscope, the animal needs to be comfortable with being held in place by the researcher.
2. Surgery 1

a. This can be either a brain or spinal cord surgery.
b. This could also be a single brain-spine surgery. However, as part of this study, the animals following a single brain-spine surgery received stimulation only under anesthesia and were imaged awake only when looking at the changes in brain-spinal activity during the transition from anesthetized to awake state. Hence, single multisite surgery animals did not follow the behavior protocol for awake study.
3. Handling and Imaging

a. Between the surgeries it is important to continue handling the animal to increase their comfort with the researcher despite the invasive procedures. A good assessment metric for determining the endpoint for handling can lie in achieving collaborative behavior qualitatively comparable to the behavior after the initial handling prior to the first surgery.
b. At this stage it can be useful to image through the implanted window to verify the ability to see the target signal. However, if not sufficient time has passed since the first surgery (e.g. at least 2 weeks for the AAV2/9-CaMKIIa-GCaMP6f viral expression used in this study), the lack of signal or responsiveness to sensory stimuli might not be informative about the prognosis for the signal strength.
4. Surgery 2

a. This can be either a brain or spinal cord surgery.
5. Habituation

a. This step consists of 3 sub-steps:
b. Habituation to the head restraint for 1-2 days, gradually increasing the duration between 5-10 minutes.
c. During the next 1-2 days, add ad libitum sucrose (30%) and brain imaging to gently promote the animal’s continuous acclimation to the parts of the experimental setup. As part of this step, continue increasing the duration of restraint towards 15-30 minutes. The microscope in this case contributes only the excitation light and it might be beneficial to avoid attempting to image during the habituation to avoid photobleaching. However, we find it useful to start imaging the brain at this earlier stage because the cost of imaging is low (data storage and experimental time), but the efforts to image at this stage help to learn the particular brain landmarks and probe the responsiveness of different areas.
d. Add hindpaw restraint and optomechanical stimulation as the final parts of the experiment. Continue increasing the duration up to 60 minutes. Continue habituating until the animal develops comfort with the needed number of trials or the animal stops improving in its performance. The performance in this case is assessed by the amount of animal’s motion and signs of not wanting to continue engaging with the experimental setup. For the purposes of sensory optogenetic and mechanical stimulation experiments (**Fig. 10**), the minimal number of trials was 60 (up to 12 minutes) to allow at least 30 trials of optical and 30 trials of mechanical stimulation.
e. As part of this study, we strongly considered the idea of habituating the animals to head-restraint during between the 1st and 2nd surgery. The benefit would be that the animal learns the behavior before any serious interventions and in theory it should be easier for the animal to learn the task after the 1st surgery than after both surgeries and re-learn the task after the 2nd surgery rather than learning it from zero after the 2nd surgery. Albeit, this approach seems promising, it has not been tested as part of this study.

**Fig. 5.**
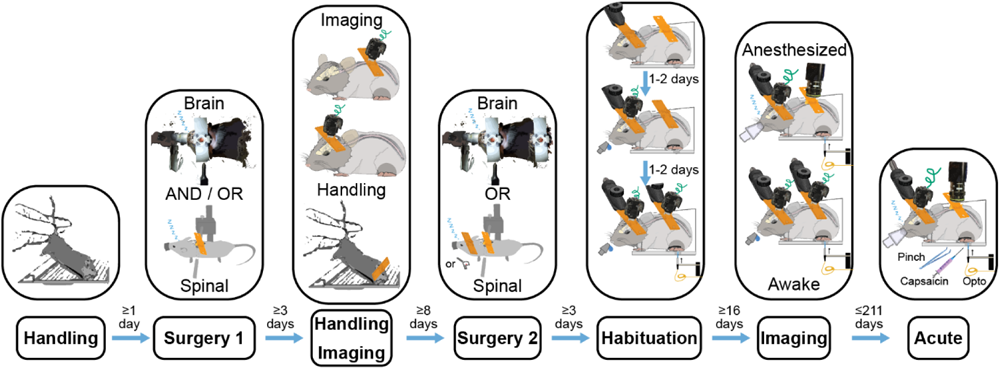
Timing of Steps that Enable Awake Simultaneous Brain-Spinal Cord Imaging. Awake imaging simultaneously in the brain and spinal cord to study sensory information processing requires careful design of a multi-step animal training protocol to maximize animal comfort and data quality.

6. Imaging

a. Imaging involves the use of the same full behavior as the last habituation step. The only difference is the addition of the microscope to image either from both sites or continue image from a single site depending on the experimental goals.
7. Acute experiment

a. As the final step, the animals typically undergo the acute experiments. In this case, every animal underwent the saline and capsaicin injections into the hindpaw to compare the impacts of the strong nociceptive stimulus on the animal’s brain-spine responses to stimulation.

Ultimately this procedure is animal-specific and all the durations provided in this procedure might have to be adjusted for a particular animal’s phenotype. In addition, given the high value of double implant animals, in the rare cases of brain implant detachment during the habituation, it is possible to reattach the implant. However, it is important to start the surgery immediately to moisturize the brain as soon as possible. The quality of the craniotomy will inevitably decrease in the case of re-implants, hence it is justified to perform this step only in the cases of animals showing exceptionally strong responses. We have not tested spinal cord re-implantation because of the lack of cases of implant detachment. However, given the propensity of the spinal cord towards bleeding and the involvement of silicone elastomer when creating a spinal cord window, the impact of spinal cord re-implantation can be predicted to be inevitably significantly more severe than in the case of brain and will likely be associated with the spinal cord injury.

Animals that were not involved in any awake behavior as part of this study (e.g. the animals participating in the spinal cord endogenous light sensitivity study) did not undergo handling, habituation and awake imaging.

### 2.6 Stimulation

#### 2.6.1 Pinching

Pinching responses were evoked by gently squeezing the hindpaw with the hand-held forceps. The effort was made to apply the pressure consistently at the same location. However, the amount of pressure exerted by every pinch varied pinch-to-pinch because of natural variability present when manually delivering somatosensory stimulation.

#### 2.6.2 Optogenetic and Mechanical Stimulation of the Hindpaw

All plantar hindpaw optogenetic stimulation experiments were performed using 470 nm light-emitting diode (LED) (M470F3, ThorLabs Inc.) controlled using an LED driver (LEDD1B, ThorLabs Inc.). Light was delivered to the plantar hindpaw using an optical fiber. The stimulated hindpaw was covered with the black aluminum foil to minimize light propagation beyond the stimulation target. The cannula of the optical fiber was glued using epoxy to the piezo-electric plate bender (CMBP09, Noliac Inc.) to allow interleaved mechanical and optical stimulation trials. The mechanical stimulation was done by poking the animal’s plantar hindpaw with the optical fiber’s cannula by actuating the piezo-electric plate bender via an amplifier. An example of the stimulation waveform parameters is shown in **Fig. 6b**.

**Fig. 6.**
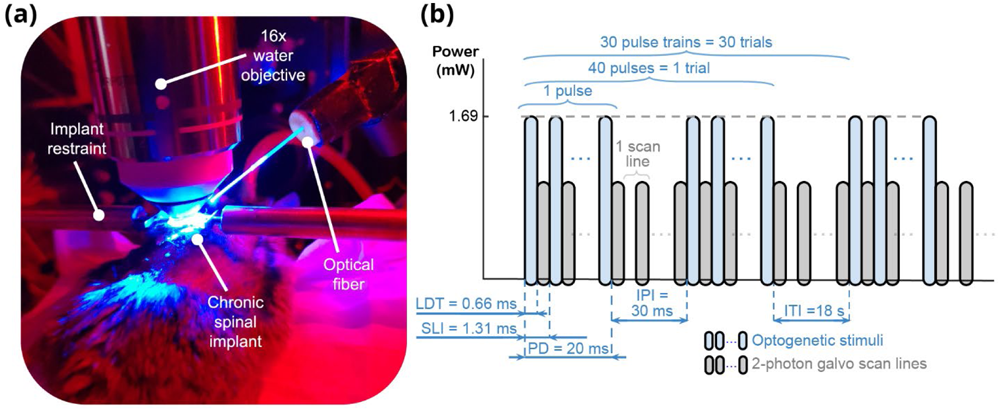
Spinal Cord Photomodulation Approach for Integratied 2-Photon Imaging. **(a)** We performed optical stimulation in the spinal cord under isoflurane anesthesia through a chronically implanted spinal cord window using an optical fiber positioned adjacent to the imaging objective. **(b)** We imaged spinal cord vascular responses using 2-photon galvo-galvo scan mode. The use of 2-photon galvo-galvo allowed us to minimize the optical stimulation artifacts by flashing the stimulation LED only during the scanner flyback between the scan lines, see Voigts et al. (2020)^57^ for further details. LDT = line dwell time, SLI = scan line interval, PD = pulse duration, IPI = inter-pulse interval, ITI = inter-trial interval.

**Fig. 7.**
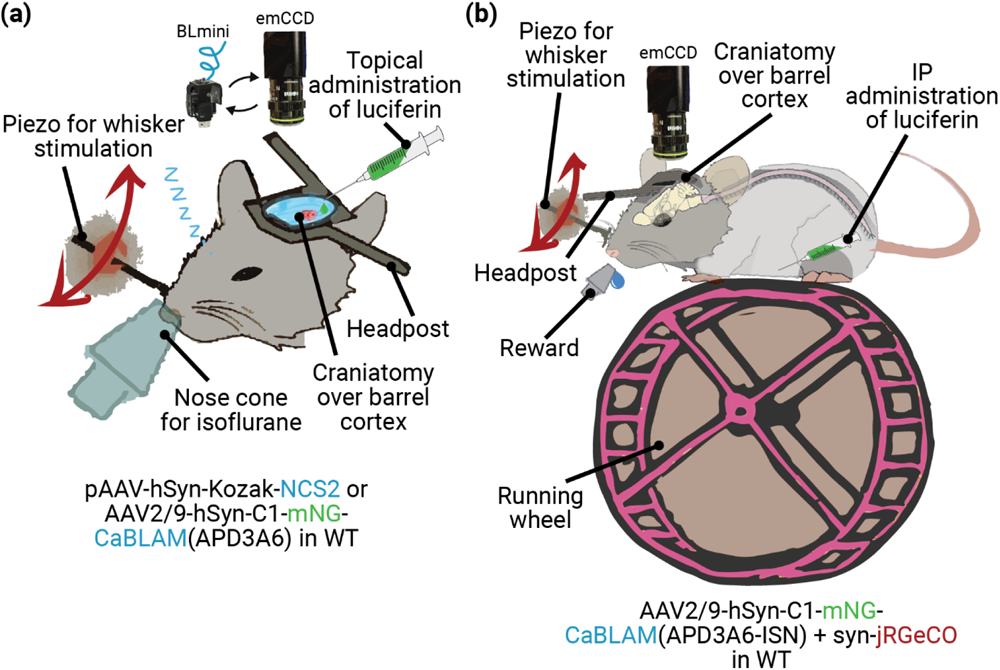
Bioluminescence (BL) Imaging Was Conducted in Anesthetized Animals With Direct Cortical Luciferin Application and In Awake Behaving Animals With Systemic Injection. **(a)** We performed acute BL imaging under isoflurane anesthesia with direct cortical administration of luciferin (Coelenterazine). The same preparation was used for imaging either using an emCCD camera or the BLmini. During the experiments testing novel BL calcium indicators (e.g., CaBLAM (APD3A6)) we stimulated vibrissae using a piezoelectric element. **(b)** For awake BL and FL (e.g., jRGeCO) imaging we administered luciferin intraperitonially (IP) before headpost-restraining the animal on a running wheel. We imaged BL and FL through the same cranial window sequentially to compare relative activation profiles.

Imaging and opto-mechanical stimulation were controlled and synchronized using either an OpenEphys or National Instruments (NI BNC-2110, NI PCI-6713 and USB-6008 for emCCD and BNC-2110 with PCIe-6353) data acquisition boards operated via MATLAB R2015b and OpenEphys software v0.5.5.3.

#### 2.6.3 Optical Stimulation of the Spinal Cord

The optogenetic stimulation of the spinal cord was performed by directly aiming an optical fiber at the spinal cord implant (**Fig. 6a**). The important aspect of spinal cord optical stimulation was the use of flyback to reduce the artifacts during 2-photon imaging. Even though the imaging approach used for collecting the data reported in **Fig. 11** relied on GFP filters that should block 470 nm light used for stimulating the spinal cord, the imperfect stop-band of the filters and high sensitivity of 2-photon photomultiplier tubes (PMTs) results in artifacts when trying to apply optical stimulation during 2-photon imaging. Flyback stimulation approach relies on the fact that galvo scanning requires pausing the imaging during the data collection with PMTs to mechanically reposition the scanner between the scan lines. The transition time between the scan lines is defined as the line period and the stimulation is applied only during this transition to effectively reduce the signal contamination with the photons reflected back into the objective from the stimulation beam.

### 2.7 Image Analysis

All microscopy images have been processed using either ImageJ (Fiji, v1.53f51) or MatLab R2021a (MathWorks Inc.). For all the multi-trial analyses (e.g. **Fig. 10, 11 and 13**) the data as plotted as a percentage change of baseline (i.e. dF/F % change) activity during 1 s interval preceding the stimulus:

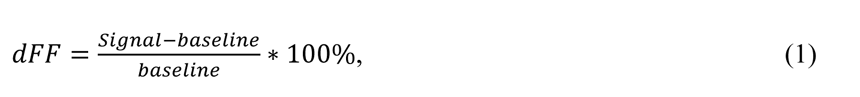

where signal is data recorded during stimulation and baseline is data recorded during the time interval preceding stimulation.

In the case of individual pinch response data in **Fig. 9**, the baseline for each neuron was defined as the full experiment-averaged activity within the area surrounding each neuron.

### 2.8 Statistical Analysis

The analysis of animal weights reported in **Fig. 8** relied on multiple comparisons using the Bonferroni method. The significance threshold was set to p-value < 0.05/300 = 0.00017. The statistical analysis was performed using MatLab R2021a (MathWorks Inc.).

**Fig. 8.**
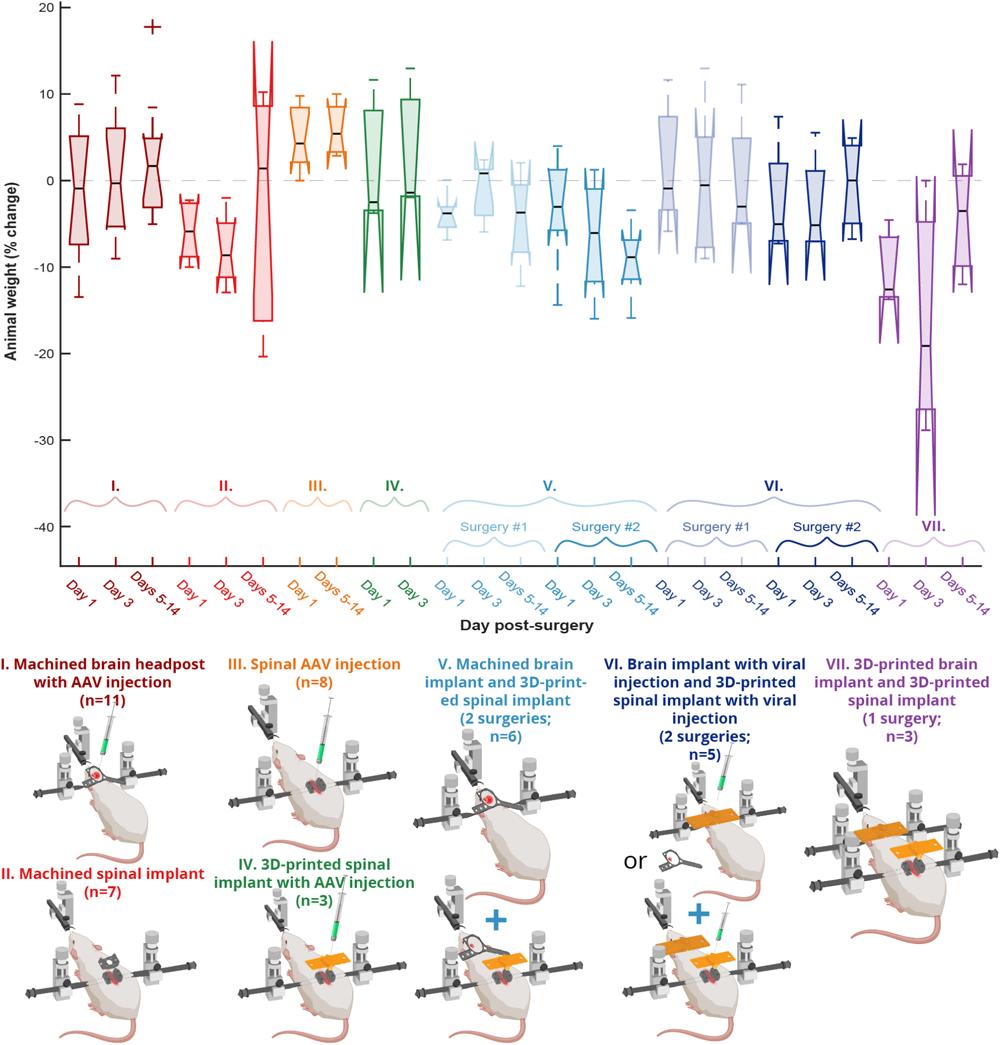
Tolerance of the Multiple Implant Variants Tested. Across implantation approaches, animals were healthy by observation, and showed either maintenance of pre-surgical body weight or recovery after ∼1-2 weeks.

## 3 Results

Here, we aimed to compare multiple implant designs and surgical preparations to drive calcium flux and thus indicator fluorescence (e.g., mechanical touch, pinching, peripheral optogenetic stimulation and local optical stimulation), as well as multiple imaging preparations (anesthetized and awake, wearable 1-photon, benchtop 1-and 2-photon imaging, sequential and simultaneous multi-organ imaging).

### 3.1 Animals Tolerate the UPRIM 3D-Printed Implants in Several Configurations

None of the implant iterations and surgical combinations tested during this work showed a significant impact on animal’s health, based on qualitative observations of post-surgical markers and behavior, and quantitative tracking of animal weight. The implants and surgical combinations tested during this work included 7 different preparations across 41 mice, summarized in Fig. 8. Our analysis identified no significant differences in weight trajectories between any of the compared groups when using a significant level adjusted for multiple comparisons (Multiple one-way ANOVAs, Bonferroni corrected, 300 surgical preparation pairs, p > .05). The surgical preparation groups were as follows:

I. Implantation of a machined brain headpost with cortical viral injections
II. Implantation of machined spinal cord implant (design described in Ref. ^17^)
III. Spinal cord viral injection without implants
IV. Spinal cord 3D-printed implant with viral injection
V. Implantation of machined brain implant (surgery #1) followed by implantation of 3D-printed spinal cord implant (surgery #2)
VI. Implantation of machined brain implant with viral injection (surgery #1) followed by implantation of 3D-printed spinal cord implant with viral injection (surgery #2)
VII. Implantation of brain implant and 3D-printed spinal cord implant in a single surgery

### 3.2 UPRIM Spinal Cord Implants are Viable for Single Cell Imaging and Have Practical Advantages

Single-site imaging of spinal cord activity dynamics has proven challenging for several reasons: (i) high mobility of the spinal cord requires sophisticated motion correction algorithms and more complex surgical implant attachment designs; (ii) imaging depth is more shallow than in the brain due to photon scattering by spinal myelination (**Fig. 2**); (iii) surgical procedures are more complex and hence require more training than brain procedures; (iv) the spinal cord has a smaller tool builder community than the brain, and advancements happen at slower pace; (v) in general the longevity of spinal cord window implants is lower than for the brain; (vi) the highly-mobile structure of the spinal cord makes it challenging to design implants cable of withstanding traditional restraint paradigms used for brain imaging, limiting the complexity and number of experimental trials that can be performed under awake spinal restraint.^58^

Several crucial breakthroughs in overcoming these challenges have been led by Schaffer and colleagues.^17,59,60^ Their spinal chamber implant design and procedure revolutionized the field by allowing chronic 2-photon imaging of blood flow in spinal stroke^59^ and axonal morphology chronic changes following spinal cord injury.^17^ Building on this success, they have now also demonstrated the feasibility of imaging ventral spinal cord using 3-photon microscopy.^60^ Their work laid the foundations for several recent studies building on this original design that have in turn made important, unique contributions.^15,58,60^

While recent progress has been substantial, prior spinal implant designs have limitations we sought to mitigate in our work. First, we found the process of restraining an awake animal using custom-fabricated fine screw-based holders time consuming and stressful for the animal when compared to the streamlined restraint process using our standard headpost implants. Second, we experienced significant variability in the quality of spinal window preparation, in part due to the need for a large amount of elastomer between the window and implant chamber.^17^ Third, we wanted to image spinal cord in awake, freely moving animals. However, the only study that has succeeded at imaging spinal cord in awake animals as we were initiating this work was done by Nimmerjahn and colleagues,^15^ and the path to adaptation to our experimental goals was unclear. Fourth, the original design reported by Schaffer and colleagues relied on potentially expensive machining and required significant time to machine in a professional workshop due to its fine dimensions and high hardness of the stainless steel required (316). We addressed these 4 constraints by the following means. First, to streamline the restraint, we adopted “restraint wings” from our headpost implant design that allow us to employ the same basic off-the-shelf clamps used for headpost restraint, and hence makes these methods easy to be adopted by any brain-focused labs. Second, to increase window longevity, we brought the glass window closer to the surface of the cord, as in our brain windows, avoiding the air gap along the midline and reducing the amount of elastomer needed. Third, to facilitate spinal cord imaging in unconstrained animals and promote broader adoption, we added a screw holder collar for the widely adopted open-source UCLA Miniscope V4. Fourth, to decrease experimental costs and streamline our ability to iterate implant designs, we used 3D-printing, manual processing (gluing, sanding, drilling and tapping by hand tools) and off-the-shelf components. Because we adopted many features for UPRIM design from brain headpost design, the final design version presented here naturally ended up being suitable for brain imaging as well. Importantly, this consistency across targets allows interchange of the same miniscope between regions imaging in the UPRIM system. While we are still testing the efficacy of our spinal implants for imaging with cellular precision, we have obtained proof-of-concept that they can allow imaging of robust single-neuron and rapid activity-specific dynamics. **Fig. 9** shows 3 neurons imaged under anesthesia using the UCLA miniscope v4 during hindpaw stimulation 37 days after implantation and AAV injection.

**Fig. 9.**
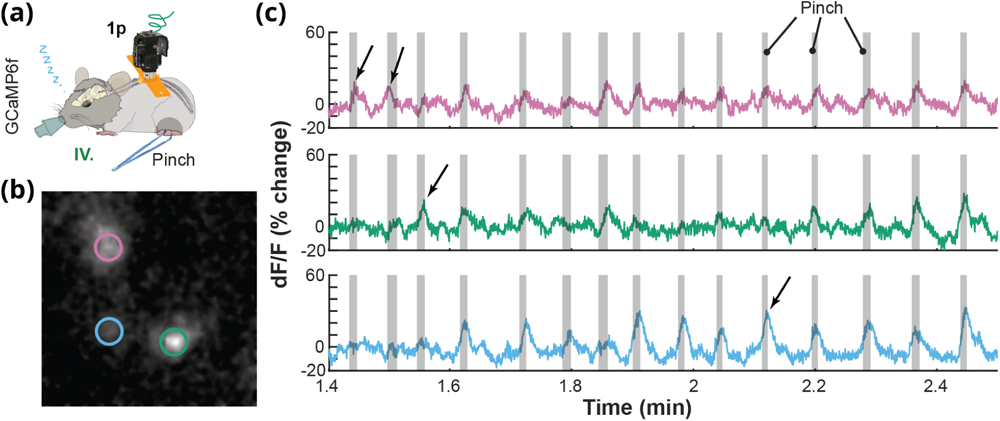
UPRIM Spinal Cord Implants are Viable for Single Cell GCaMP Imaging. **(a)** Individual cell GCaMP6f calcium responses to hind paw pinch were imaged under anesthesia in an animal implanted with spinal cord UPRIM via surgical procedure # IV and using Miniscope v4. Calcium indicator was expressed via AAV2/9-CaMKIIa-GCaMP6f-P2A-nls-dTomato. **(b)** A pinch response map as the average of multiple trials, showing activation in multiple distinct neurons (pink, green and blue circles). **(c)** Time series show changes in signal for each color-matched cell. Grey shadings highlight the pinching time intervals, arrows emphasize responses that differed between 3 cells.

Our initial imaging success (and see further examples throughout) is paired with a variety of other pragmatic benefits of UPRIM as a spinal implant system. First, as shown in **Fig. 8**, these implants are well-tolerated, even when multiple windows are in place. We observed no specific impact on body weight. Second, as described throughout, the UPRIM system is lightweight, at only 1.64 g compared to 1.96 g for the stainless still machined implant reported by Schaffer and colleagues (this weight includes the chamber, glass window, screws and spinal bars). In the case of brain imaging, UPRIM weighs only 1.45 g whereas commonly employed machined versions weigh 2.39 g (including restraint chamber and glass window). Altogether, a pair of UPRIM multi-organ implants would weigh 3.09 g, versus 4.35 g for machined implants, a 29% reduction. This UPRIM value includes the additional features necessary for miniscope attachment, which if added to the machined implants would increase their weight differential. Third, the UPRIM fabrication process is easier than custom machining, and because it does not require a professional machinist is also cheaper.

### 3.3 Imaging Calcium Activity in the Brain and Spinal Cord with UPRIM

Single brain-spinal cord surgery allows chronic imaging of transgenic GCaMP6f responses for up to 15 days post-surgery (**Fig. 10a**). Dynamic 1-photon calcium responses of transgenically labeled somatostatin (SOM) interneurons were reliably localized to the side of stimulation for up to 154 days after the surgery. We also conducted sequential imaging from brain and spinal cord UPRIM under anesthesia (**Fig. 10b**) and simultaneously in awake animals (data not shown). In the example shown in **Fig. 10b**, imaging of nociceptive responses following optogenetic stimulation in Trpv1^Chr2EYFP+/−^ mice was first performed in the brain 24 days after surgical implantation with UPRIM. After 52 days, an additional UPRIM was implanted in the spinal cord, and the data shown acquired 76 days later. Plantar optical stimulation of the right hindpaw reliably evoked cortical responses to all stimuli except for the weakest stimulation (i.e. 0.28 mW). Ipsilateral plantar stimulation of the left hindpaw also evoked brain responses, but of smaller (1% change vs. 2.5% change for the strongest optical stimulus) and more sustained magnitude compared to the contralateral stimulation. In contrast, the spinal cord responded to mechanical plantar stimulation of the left hindpaw, with no detectable response to right hindpaw plantar stimulation.

**Fig. 10.**
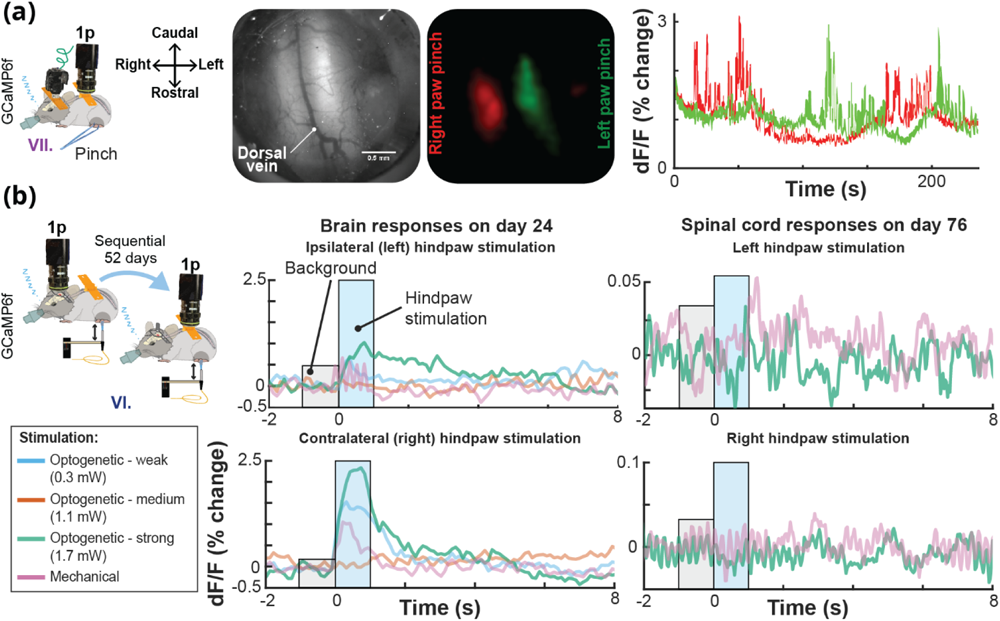
Calcium Activity in the Brain and Spinal Cord in the Same Mouse Using UPRIM. **(a)** Lateralized localization of responses to hindpaw pinch in the spinal cord using the UPRIM implant and 4x magnification benchtop imaging. The GCaMP6f indicator was expressed in Somatostatin-positive interneurons (SOM). Images in this panel from left to right show: The experimental setup with spinal cord reference axes for orientation, spinal cord brightfield image, GCaMP6f average response maps, and time courses showing the independence of relative light intensity signals for left (*green*) and right (*red*) cord regions. In this experiment, the mouse was implanted with a brain and spinal cord UPRIM (surgical procedure # VII). **(b)** UPRIM calcium responses in cortex to optogenetic and mechanical stimulation 24 days after implantation, and in the same mouse in the spinal cord 52 days after implanting and imaging brain UPRIM (surgical procedure # VI). Both data sets were acquired under anesthesia. This was a Trpv1^ChR2eYFP+/-^ animal injected with a mixture of two non-flex AAVs to co-express GCaMP6f and jRGeCO both in the brain and spinal cord.

### 3.4 Using UPRIM to Test Optical Activation of the Spinal Cord: Vasodilation of Spinal Vessels in response to Blue Light in Wild-Type Mice

Multiple studies have shown that blue light can, through an endogenous mechanism, drive dilation.^61,62^ Using UPRIM, we tested that blue light (470 nm) induced vessel dilations in wild-type animals using 2-photon imaging 24 days after spinal surgery (**Fig. 11**). We found reliable induction of vasodilation in several, but not all, vessels tested.

**Fig. 11.**
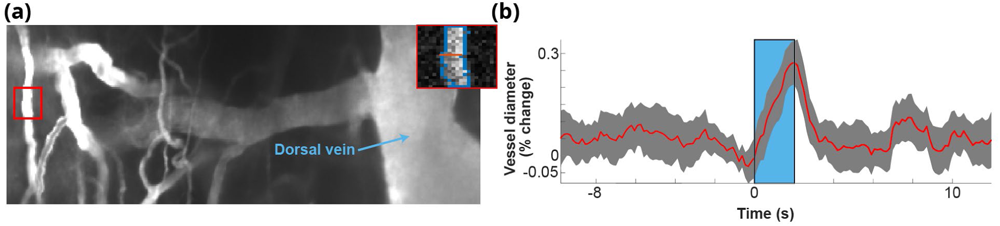
Vascular Dilations in the Spinal Cord in Wild Type Mice Driven by 2 sec Long Trains of Pulsed Light. **(a)** Map of spinal cord vasculature obtained with 2-photon imaging after intravenous 10 kD FITC-green dextran injection. Zoomed-in vessel inset in top right shows a single frame of a segmented vessel during stimulation. **(b)** Example of a responsive vessel segment.

### 3.5 Bioluminescence Offers Benefits for Wearable Miniscopes: Smaller, Fewer Component, and Lower Power Consumption

Based on the advantages of BL imaging in general, and several benefits to miniscope design conceptually possible, we developed a novel miniscope optimized for BL imaging, the ‘BLmini.’ **Fig. 12** summarizes characteristics of two generations of FL and BL-targeted miniscopes. The signal strength comparison between FL v3.2 and BLmini v1.0 miniscopes was determined by measuring minimal detectable optical signal with two microscope configurations. The relative signal strength for FL v4 and BL v2.0 miniscopes was based on based on 5x increase in sensitivity of miniscope v4 compared to v3.2 shared with us by the miniscope developers. The power budget in **Fig. 12** was estimated for a wire-free version of UCLA miniscope made public on 5/21/2019.

**Fig. 12.**
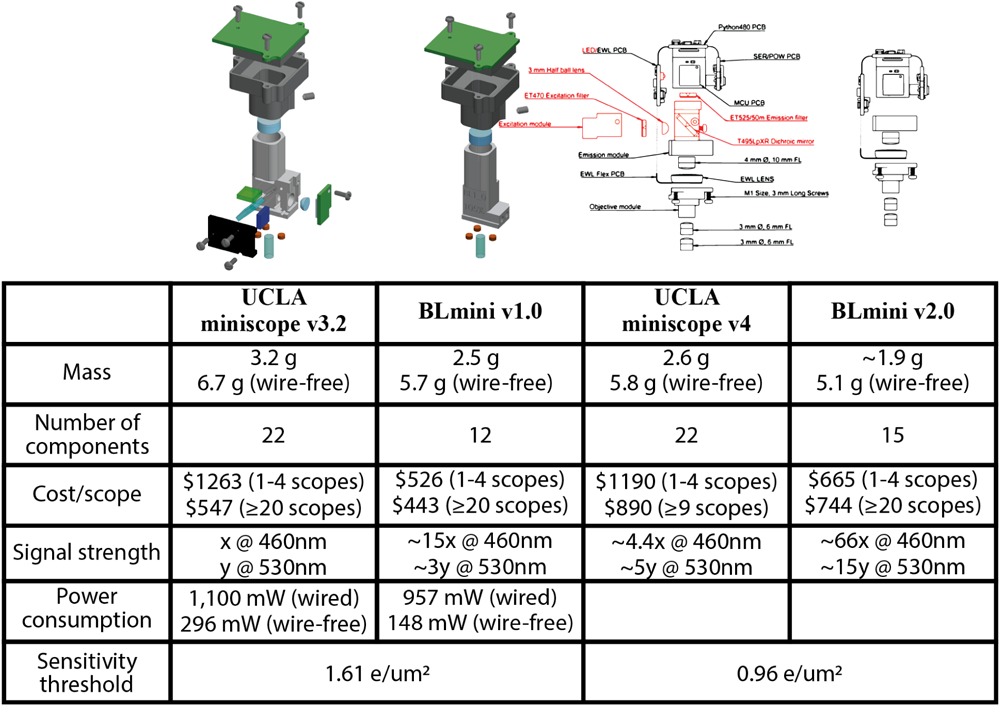
Bioluminescent Imaging Requires Fewer System Components, Decreasing Miniscope Complexity and Weight. Conversion of UCLA miniscopes v3.2 or v4 from FL-centric to BL-centric design offers 22-27% reduction in mass, 32-45% reduction in the number of components and assembly labor, 19-58% reduction in cost/scope, 3-66 times higher sensitivity for optical signals that do not require filtering, 13-50% reduction in electrical power consumption.

### 3.6 Using Miniscopes to Detect Bioluminescent Indicators

BLmini v1.0 and v2.0 were able to capture temporal dynamics of *in vivo* BL that tracked those observed using an electron-multiplying gain camera (emCCD Ixon 888, Andor Inc.) (**Fig. 13a**). For comparison, we overlaid time courses acquired using the BLmini v1.0, BLmini v2.0 and emCCD Ixon 888, each from a different experiment (Ref. ^63^; **Fig. 13a**). The signal captured by the BLmini also tracked more subtle features of the known BL response. These properties include an early slowing in photon production during substrate injection, thought to be due to the inhibition of photon production by the breakdown products.

A difference between emCCD and miniscope signals is their baseline signal magnitude prior to the injection of h-CTZ. The CMOS sensor employed in the miniscope does not rely on any form of cooling and consequently suffers from significantly stronger thermoelectric background noise, as well as anisotropic distribution of noise across the pixels. The most critical factor that helped us to mitigate these issues involved letting the miniscope to operate for 30 minutes (duration was determined empirically) before starting to image. This wait time does not reduce the noise, however, it offers a stable baseline for measurements.

**Fig. 13.**
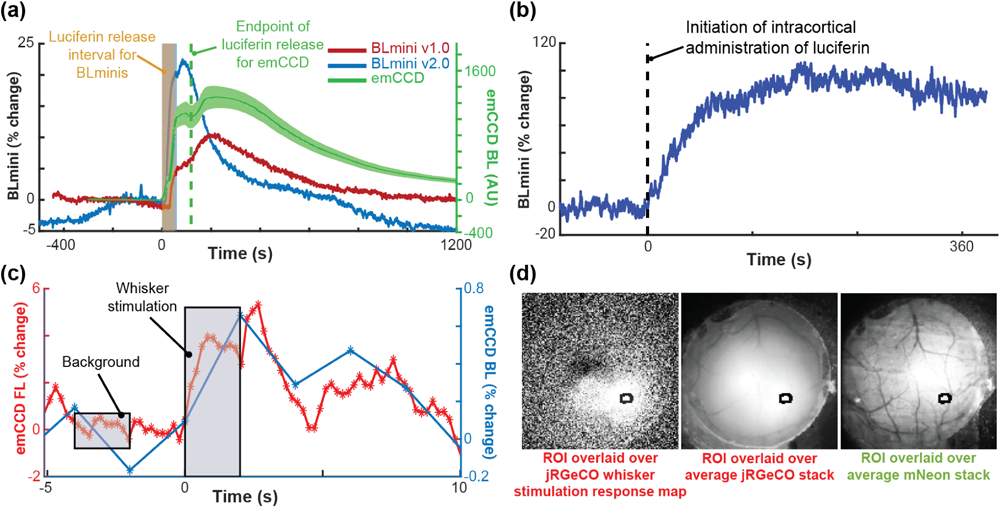
Imaging Bioluminescent and Fluorescent Indicators Using the BLmini and FL. **(a)** Time courses of BL photon production detected by an emCCD (ANDOR) and two generations of BLmini miniscopes in three different mice. Luciferin (CTZ) was directly applied to the cortical surface, previously transduced with either LMO3^63^ (emCCD data) or NCS2^65^ (BLmini data). **(b)** BLmini v2.0 allows detection of photon production by the calcium indicator CaBLAM (APD3A6).^55^ **(c)** Time series for BL and FL calcium indicator output from a single imaging session, from CaBLAM (APD3A6-ISN) and jRGeCO (with full field excitation illumination) virally transduced in SI Vibrissal Barrel Neocortex (synapsin (Syn) promoter). In these data, vibrissal stimulation was applied during either BL (blue) or FL (green) imaging (25 trials in each condition). **(d)** The Region-of-Interest used to extract jRGeCO and CaBLAM (APD3A6-ISN) signals overlaid on the mean jRGeCO vibrissa response map (left), the mean red FL intensity map across all frames (center) and the mean green FL map, reflecting expression of mNeon-green (mNG) in the CaBLAM (APD3A6-ISN) construct.

Using the more sensitive BLmini v2.0, we were able to measure not only co-factor independent trajectory of light emission, but also calcium-dependent activity (**Fig. 13b**). Further, BL indicators themselves are reaching the stage of development where we can image using emCCD camera sensory stimulation responses of comparable time course using BL and FL in the same preparation (**Fig. 13c and 13d**).^55^

## 4 Discussion

The hardware and surgical advances we present here enabled observing ‘field’ calcium and vascular dynamics in brain and spinal cord, and from the same animal for up to 8 months (data not shown) after the first surgery. The novel design provides a well-tolerated, affordable and common format for imaging the brain and spinal cord, in fixed and free-behaving mice. As such, this approach enables unique observations under conditions where each site is imaged individually but in the same mouse and on the same day, such as the time course of plastic changes in neural ensembles following a peripheral manipulation. The UPRIM strategy also enables unique correlational studies with simultaneous imaging across sites, for example in determining conjoint spinal cord and brain activity that may be key to nocifensive responses or detection of nociceptive stimuli.^64^

The main limiting factor for imaging individual cells across multiple sites was the longevity of the spinal cord window due to overgrowth. Our maximal duration for obtaining quality signals was 37 days after the spinal cord surgery. This finding aligns well with previous observations using stainless steel implants that resolved spinal cord axons for more than 35 days in only ∼50% of implanted mice due to fibrosis.^17^ Maintaining good surgical technique remains the best tool to increase the longevity of the spinal cord windows. In our experience, the key surgical priority is minimizing injury and bleeding. Other important factors leading to reduced insult to the spinal cord include: (i) sealing all bone edges throughout the laminectomy using tissue glue (e.g., VetBond); (ii) minimizing elastomer use (e.g., KwikSil; in the implant design we present here, this was achieved by positioning the glass window under the implant chamber, akin to traditional brain implant designs); (iii) increasing the pressure seal between the glass window and the spinal cord (akin to traditional brain implant designs); and, (iv) using a pneumatic drill rather than a mechanical drill or scissors. Alternatively, reducing surgical invasiveness by imaging between the vertebrae prolongs the chronicity of spinal cord imaging for up to 167 days.^66^ Also, treating an implant with gradually releasing anti-inflammatory agents might increase the duration of single-cell resolved imaging by suppressing fibrosis.^58^ Besides surgical preparation, habituation of the animal remains an equally important factor in maintaining spinal cord window quality when it comes to awake behavior. We found that high mobility of the spinal cord makes it more prone to injury. Even in the instantiation of miniscope imaging in freely behaving animals, short-term (∼2-3 minutes) restraint is necessary to attach the miniscope. We observed that even 2-3 minutes of restraint disturbs the quality of the spinal cord window in a poorly habituated mouse. Therefore, taking extra care in habituating to restraint and handling is essential. Alternatively, anesthetizing the animal allows simplified attachment of the miniscope. In our experience, attaching the miniscope under anesthesia works well for imaging unrestrained animals, but cannot be easily applied to restraint experiments because animal awakening can cause sudden and forceful movements that damage the implant. However, every cycle of anesthetic use (e.g., of isoflurane) can have an additive negative impact on the animal’s health and ultimately reduces study duration.^27^ We summarize the overarching timeline describing different major procedures and time intervals for each procedure as implemented for the final awake behavior experiments in this study in **Fig. 5**.

Similar to the spinal cord implant design described in Ref. ^67^, our use of plastic for the implant chamber fabrication makes it more compatible with photoacoustic, ultrasound, and MRI imaging, as well as ultrasound stimulation and radiation therapy. We have not tested non-optical methods as part of this development process, and additional modifications might be needed to avoid artifacts. However, for non-miniscope imaging, set-screws are not necessary, and the plastics we tested can be threaded, decreasing component number. An important factor to consider when threading plastics is that these threads are more fragile than those in affixed nuts. In our experience, tapped threads reliably serve ∼2-3 screw-unscrew cycles. Replacing 000-120 screws with glue could be a solution, but requires additional testing and we have not yet validated this approach.

## 5 Limitations

We have not applied the 3D-printed implant design described here for BL imaging. However, the metal implants we used for BL imaging (**Fig. 13**) have the same optical properties as 3D-printed implants we used for FL imaging (**Fig. 10**). Both implant designs rely on identical glass window-tissue interfaces, and hence there is no reason to expect signal differences between them. We were able to image calcium responses of individual neurons in the spinal cord for up to 37 days and dynamic ‘field’ 1-photon responses for up to 154 days. The dominant form of spinal cord implant failure was overgrowth of the window. As this report was in final stages of preparation, the first progress in mitigating the fibrosis in spinal cord optical windows was described in Ref. ^58^. Deeper assessment of animal immune responses and more prolonged chronic imaging in awake behaving animals across both brain and spinal cord, especially using microscopes optimized for BL, are key next steps.

## 6 Conclusion

Here, we successfully developed and implemented a novel 3D printed brain-spinal cord implant, and advanced/refined surgical and sensory modulation procedures, to achieve simultaneous miniscope and two-photon imaging of the brain and spinal cord. This progress overcomes previous limitations, enabling detailed and prolonged observation of neural and vascular dynamics across multiple organs. Our integration of BL imaging, with its novel indicators and modified miniscopes (BLmini), offers a potential alternative that can enhance multi-organ studies, offering simplified microscope design without confounding photoeffects. These advancements open new pathways towards developing deeper understanding of neural processes, especially in sensory behaviors and pain perception.

## Code, Data, and Materials Availability

The design files, codes and data are available through the following DOI: https://doi.org/10.26300/2yeh-tr48.

## Acknowledgments

This work was supported by grants from the National Science Foundation (NSF NeuroNex 1707352) and the National Institute of Health (NIH NINDS R01NS108414, and U01NS099709). Authors would like to thank Ute Hochgeschwender for providing AAV constructs, luciferin samples and valuable discussions of experimental design of bioluminescence experiments. We thank Daniel Aharoni for support and valuable guidance in working with miniscopes, Chris Schaffer and his team for providing a tutorial on spinal cord surgeries, and helpful discussion on imaging and designs of the spinal window implants. We would like to thank Thomas Kirkland and Joel Walker for providing fluorofurimazine samples for awake BL imaging. We also thank Kimani C. Toussaint Jr. for valuable discussions around empirical testing of miniscopes, Adriana C. Salazar Coariti for valuable discussions around experimental design and visualization, Jill Juneau for valuable discussions about electrical circuit design and troubleshooting, and Anusha Allawala for contributing to initial establishment of spinal cord surgeries and immunohistochemistry. We thank all members of the Bioluminescence Hub (www.bioluminescencehub.org) for helpful discussions, especially Dr. Justine Allen for myriad levels of programmatic supporting and coordinating collaborative efforts, and Laurie Lynch for animal colony and equipment management.

The following figures were created partially or fully with BioRender.com: Fig. 1, Fig. 8 and Fig. 14a. Fig. 2, 12 and 13 were modified, with permission, from Ref. ^65^.

**Fig. 14.**
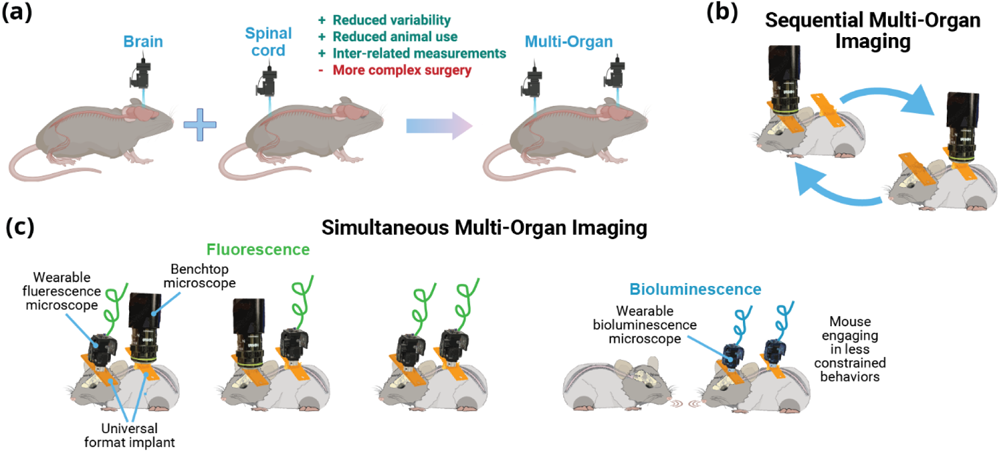
Beyond Brain-Only Imaging Offers Many Opportunities for Experimental Designs and Hardware Combinations (a) In this report we demonstrate feasibility of gaining chronic optical access to the brain and spinal cord within the same animal. Universal format implants described as part of this report allow sequential **(b)** and simultaneous **(c)** imaging of the brain and spinal cord within the same animal using benchtop and wearable microscopes via FL or BL. Wearable microscopes for BL imaging, and hence the implants, can be designed smaller, allowing animals to engage in more natural behaviors.

Author contributions: D.C. Conceptualization, Data Curation, Formal Analysis, Investigation, Methodology, Visualization, Writing – Original Draft Preparation, Writing – Review & Editing. C.B. Resources. J.M. Resources. A.B-A. Investigation. N.F. Investigation. N.S. Resources, Conceptualization, Funding Acquisition. C.S. Funding Acquisition. M.G-R. Resources. D.L. Resources, Writing – Review & Editing. D.A.B. Conceptualization, Funding Acquisition, Resources, Supervision, Writing – Review. C.I.M. Conceptualization, Funding Acquisition, Resources, Supervision, Writing – Review & Editing.

**Dmitrijs Celinskis** serves as a Postdoctoral Research Associate at the Carney Institute for Brain Science at Brown University. His area of specialization lies in biomedical engineering, focusing primarily on neurophotonics, along with the design and fabrication of devices. He is adept in both benchtop and in vivo testing, as well as in analyzing 3D time series data. Dr. Celinskis brings a wealth of knowledge in neuromodulation, medical imaging, and neurobehavioral methods, making him an indispensable contributor to cutting-edge neurotechnology initiatives.

**David A. Borton,** an Associate Professor of Engineering and Brain Science at Brown University, Associate Professor of Neurosurgery at the Rhode Island Hospital, and a Biomedical Engineer at the Providence VA Medical Center’s Center for Neurorestoration and Neurotechnology. David Borton received his B.S. in Biomedical Engineering from Washington University in St. Louis in 2006, his PhD in Bioengineering from Brown University in 2012, and performed a Marie Curie post-doctoral fellowship at the Ecole Polytechnique Fèdèrale de Lausanne from 2012 to 2014. Prof. Borton leads an interdisciplinary team of researchers focused on the design, development, and deployment of novel neural recording and stimulation technologies.

**Christopher I. Moore**, an Associate Professor at the Brown Institute for Brain Sciences, specializes in systems neuroscience. He earned his Ph.D. from MIT and held a postdoctoral fellowship at Harvard Medical School. Dr. Moore’s notable contributions include work on thalamic reticular nucleus and its role in neocortical spindles, highlighted in Nature Neuroscience. His interdisciplinary approach, combining neuroscience and philosophy, has been recognized with numerous honors, including the Angus N. MacDonald Excellence in Teaching Award at MIT.

